# The effects of elevated serum lactate on measures of apparent cerebral metabolism and perfusion in the anesthetised rodent brain

**DOI:** 10.64898/2025.12.17.695057

**Authors:** Daniel Radford-Smith, Catriona Rooney, Jordan McGing, Ayaka Shinozaki, Surrin S Deen, Fulvio Zaccagna, Richard Healicon, Vicky Ball, Daniel Anthony, Damian J Tyler, James T Grist

## Abstract

Cerebral metabolism is coupled to perfusion and can be probed using a combination of hyperpolarized carbon-13 MRI and perfusion weighted imaging. It is unknown how alterations in serum lactate affect apparent lactate exchange and pyruvate dehydrogenase flux using hyperpolarized ^13^C MRI, and so this pre-clinical study was designed to assess the sensitivity of MRI to detect changes in perfusion and apparent metabolism in the anesthetized rodent brain due to sodium lactate administration.

Results demonstrated significant differences in cerebral blood flow between the saline and sodium L-lactate administered at 0.5 g/kg (112 ± 5 vs 127 ± 2 mL/100g/min), saline and sodium L-lactate administered at 1 g/kg (112 ± 5 vs 90 ± 5 mL/100g/min), and between sodium L-lactate at 0.5 g/kg and sodium L-lactate at 1 g/kg (127 ± 2 vs 90 ± 5 mL/100g/min).

There was excellent reproducibility of cerebral perfusion measurements (ICC > 0.95) between observers, indicating high reliability in these results. There were significant differences in [1-^13^C] lactate:[1-^13^C] pyruvate between saline vs 0.5 g/kg (0.22 ± 0.05 vs 0.59 ± 0.15) and saline vs 1 g/kg (0.22 ± 0.05 vs 0.62 ± 0.04), but no significant differences in ^13^C bicarbonate:[1-^13^C] pyruvate (p = 0.1).

In conclusion, perfusion and [1-^13^C] lactate:[1-^13^C] pyruvate are sensitive to changes in serum lactate, but ^13^C bicarbonate:[1-^13^C] pyruvate were not. This has implications for brain studies using [1-^13^C] pyruvate, and understanding the impact of serum lactate on clinical hyperpolarised imaging is needed.

## Introduction

Cerebral metabolism is coupled to cerebral blood flow, a process which is thought to involve a complex interplay between neurons and astrocytes that is critical to normal brain function^1^. For example, the delivery and uptake of glucose in neurons may be regulated by astrocyte mediated vasoconstriction and dilation, and the shuttling of lactate from astrocytes to neurons fuels neuronal mitochondria^2,3^. However, the coupling of cerebral metabolism to blood flow also needs to be able to accommodate other brain energy demands that may arise during injury or disease. For example the activation of microglia from a quiescent to an activated state demands an increase in the rate of glycolysis^4^. Indeed, neuropathology that results in alterations in cerebral perfusion and apparent metabolism is often measured in the clinic using ^18^F-FluroDeoxyGlucose Positron Emission Tomography (FDG-PET), which is a biomarker of glucose perfusion and uptake^5,6^. However, FDG uptake can be confounded by alterations in the haemodynamic response, nutrient intake and blood chemistry, and the direct detection of mitochondrial metabolism has remained beyond the reach of conventional clinical imaging methods^7^. Indirect methods that probe perfusion are also commonly used as a surrogate for cerebral metabolism^8,9^. The combination of techniques that directly report on cerebral metabolism and perfusion coupling *in vivo* overcomes some of the limitations of the methods when they are used independently. Gadolinium based perfusion MRI employs imaging to track the transit of a gadolinium-containing contrast agent to estimate cerebral perfusion and blood volume and hyperpolarized carbon-13 MRI (HP ^13^C MRI) allows for the real-time detection of lactate exchange and Pyruvate Dehydrogenase (PDH) flux ^10–15^. However, it remains unclear how perturbations of blood chemistry might impact the interpretation of results from combined ^13^C/perfusion studies of the brain.

L-lactate is known to be a signalling molecule and substrate for cerebral metabolism^16,17^. Previous studies in the diabetic brain have shown that low dose lactate administration can recover Cerebral Blood Flow (CBF), and lactate has been used as a promising therapeutic agent post-reperfusion to reduce lesion volume post-stroke in rodents^18,19^. Despite this, the effect of raising blood lactate levels on apparent cerebral metabolism and perfusion coupling *in vivo* has not been explored in pre-clinical studies. This study aimed to assess the variability in measures of apparent [1-^13^C] pyruvate metabolism and perfusion in the anesthetised rodent brain when challenged with sodium L-lactate.

## Methods

All animal experiments were carried out under UK Home Office regulations. 13 male Sprague Dawley (SD) rats were purchased from Charles River (200-220g at purchase) and housed in a 12-hour light/dark cycle with free access to food and water.

Animals were imaged using a 60:40% O_2_:N_2_O protocol. 10 SDs were imaged to estimate baseline cerebral perfusion. Animals were anesthetised with 2.5% isoflurane, a tail vein cannula was inserted, and a line drawn across the mid brain using permanent marker to ensure reproducible positioning. 3 SDs were imaged three times in a cross over study, and each received a 0.5 mL intra-peritoneal injection of saline or sodium L-lactate (two doses used on different days, 0.5 and 1 g/kg) approximately 10 minutes prior to ^13^C spectroscopy. Animals were reduced to 2% isoflurane when in the scanner (7T, Agilent, Santa Clara, USA). Serum lactate was measured from a tail vein before lactate injection and at the end of scanning using a hand-held lactate monitor (EKF diagnostics, Barleben, Germany).

### Hyperpolarization

Approximately 30 mg samples of [1-^13^C] pyruvic acid (Merck, New Jersey, USA) were mixed with the trityl radical, OX063 (15 mM) (Oxford Instruments, Abingdon, UK) and gadolinium (3 μL, 1:50 dilution in water) (Dotarem, Guerbet, France) and were hyperpolarized in a prototype system at 3.35 T for approximately 1 hour. Samples were dissolved using sodium hydroxide buffer (approximately 4.5 mL) and 1 mL of the final solution was injected over 6 seconds with a 200 uL saline chase.

### MRI

For ^13^C spectroscopy, animals were placed in a custom-built cradle on a 2-channel ^13^C receive coil (Rapid Biomedical, Rimpar) using a ^1^H/^13^C volume coil for radiofrequency transmission (Rapid Biomedical, Rimpar). Respiratory monitoring was performed throughout each experiment, targeting 60 breaths per minute. Localiser images were acquired and a pulse-acquire spectroscopic slab placed through the brain of each animal (gaussian excitation pulse, excitation pulse width = 100 us, receive bandwidth = 5 kHz, TR = 1 s, TE = 0.3□ms, slice thickness = 20 mm, FA = 15 degrees, number of time points = 240). Acquisition of data was started prior to injection.

After the ^13^C injection, animals were transferred to a second custom cradle with a tooth bar and ear pins, using a 4-channel ^1^H phased array head coil (Rapid Biomedical, Rimpar), for perfusion (200μL Dotarem with 200μL saline chase, single slice 2D Gradient Echo, slice through mid-brain, TR = 20 ms, TE = 10 ms, FA = 20 degrees, acquisition matrix = 128×64, FOV = 35 mm, slice thickness = 1 mm, total number of time points = 60, time resolution = 1 s, 3 baseline images pre gadolinium-injection) and anatomical T_1_-weighted imaging pre-and post-gadolinium bolus (Repetition time (TR) = 5 ms, Echo time (TE) = 1 ms, Flip angle (FA) = 12°, RF spoiled, acquisition matrix = 256×256×64, reconstruction matrix = 512×512×64, field of view (FOV)= 40×40×40 mm^3^).

### Post-processing

Summed spectra were processed using jMRUI (v5.1) using [1-^13^C] pyruvate, [1-^13^C] lactate, ^13^C bicarbonate, [1-^13^C] alanine, and [1-^13^C] pyruvate-hydrate as basis functions. The [1-^13^C] lactate:[1-^13^C] pyruvate, ^13^C bicarbonate:[1-^13^C] pyruvate and ^13^C bicarbonate:[1-^13^C] lactate ratios were calculated for each acquired data set.

Perfusion data were post-processed using model-free deconvolution^20^. A region of interest was drawn in the supplying vessels forming Cerebral Blood Flow (CBF), Cerebral Blood Volume (CBV) – relative to the Arterial Input Function (AIF), Time to peak (TTP), and Mean Transit Time (MTT) maps.

### Image analysis

To assess the reproducibility of the analysis of the perfusion data, analysis of the No-Intervention animals was performed by 5 observers who independently drew regions of interest in the whole brain taking care to exclude the supplying vessels using T_1_ weighted imaging pre-and post-contrast for guidance. ROIs were drawn on the CBF map, and then transferred to CBV, TTP, and MTT images.

For the 3 rats that also received a [1-^13^C] pyruvate injection, one observer drew a whole brain region of interest, taking care to exclude the supplying vessels, as described above.

### Statistical analysis

All statistical analysis was performed in R (4.0.5). The intra-class correlation coefficient (ICC) was calculated for inter-observer reliability of perfusion measurements, a cut-off of <0.5 was assumed to show poor reliability, 0.5 to 0.75 to indicate moderate reliability, 0.75 to 0.9 indicating good reliability, and >0.9 indicating excellent reproducibility ^21^.

A Kruskal-Wallis test was used to assess variance between the treated animals. If a significant contribution to variance was detected, then paired post-hoc tests were performed between groups using a Bonferroni correction for multiple comparisons.

Correlations between perfusion and ^13^C metabolic measures were assessed using Spearman’s correlation. A p-value of < 0.05 was used as a threshold for statistical significance.

## Results

### Reproducibility of cerebral perfusion measurements in the healthy rodent brain

With multiple observers drawing ROIs in the brain of the rats. There was excellent reproducibility for CBF measurements (0.96 [0.92-0.99], lower bound – upper bound, respectively), CBV (0.98 [0.97-0.99], lower bound – upper bound, respectively), TTP (0.98 [0.96-0.99], lower bound – upper bound, respectively), and MTT (0.99 [0.98-0.99], lower bound – upper bound, respectively).

### Lactate modulates cerebral perfusion in the anesthetised rodent brain

Example, CBF maps and hyperpolarized spectra from cross-over study are shown in Figure 1 A-C and D-F, respectively.

**Figure 1.**
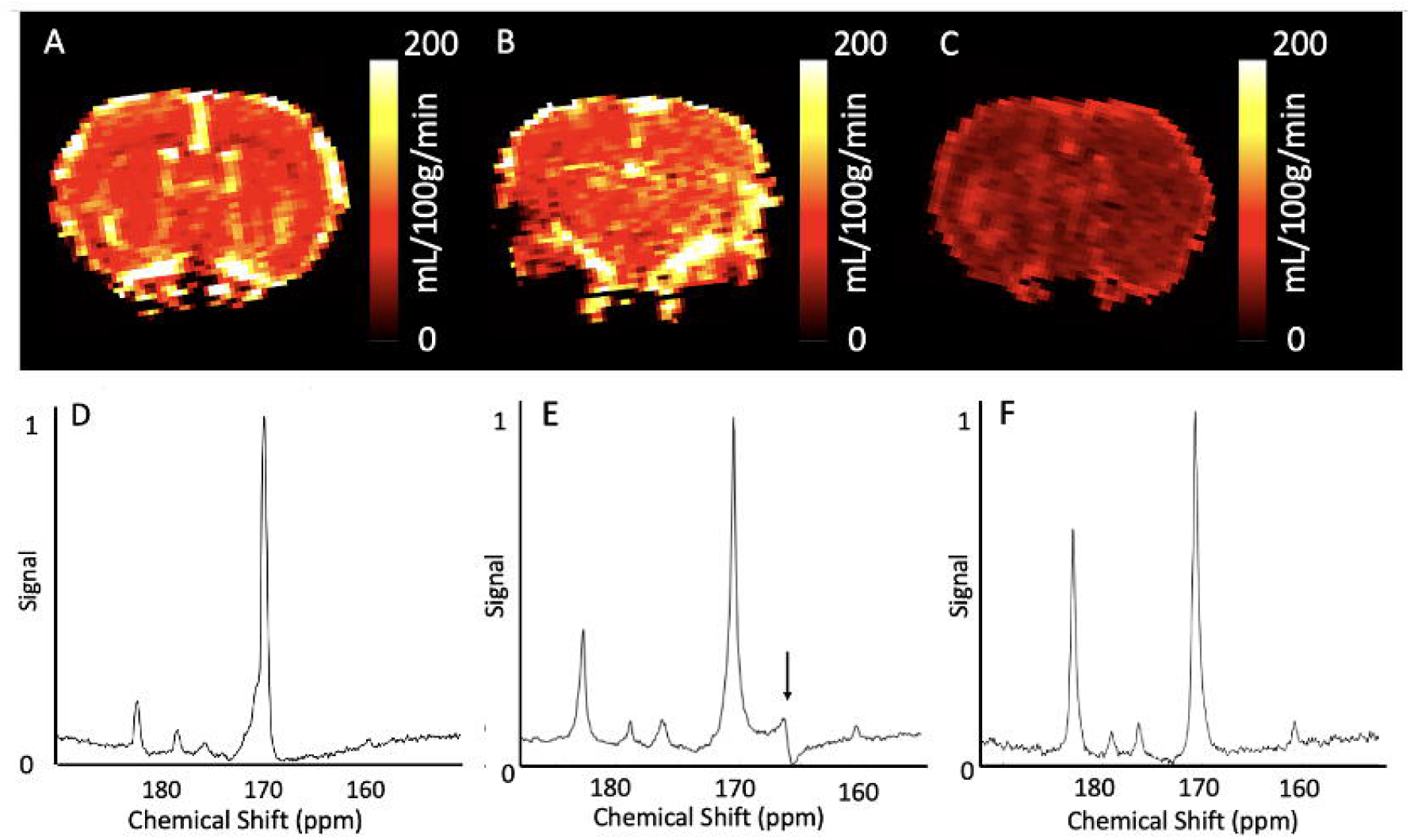
Example calculated Cerebral Blood Flow maps after an I.P injection of saline (A), half-dose (B), and full dose (C) lactate from the same animal. Hyperpolarized [1-13C]pyruvate spectra after I.P. injection of saline (D), half-dose (B) and full-dose (C) lactate from the same animal. Signal from a ^13^C Urea standard is seen in E, denoted by a black arrow.

It was found that there were significant differences observed in CBF between saline and 0.5 g/kg lactate (112 ± 5 vs 127 ± 2 mL/100g/min), saline and 1 g/kg lactate (112 ± 5 vs 90 ± 5 mL/100g/min), and 0.5 g/kg and 1 g/kg lactate (127 ± 2 vs 90 ± 5 mL/100g/min) groups. CBV, however, did not significantly change with lactate dose – with the saline (0.74 ± 0.06 mL/100g), 0.5 g/kg lactate (0.97 ± 0.19 mL/100g), and 1 g/kg lactate (0.88 ± 0.1 mL/100g) groups showing similar CBV values (Group-wise p = 0.6).

MTT varied significantly between the saline and 1 g/kg lactate (0.39 ± 0.05 vs 0.62 ± 0.04 s^- 1^) and the 0.5 g/kg and 1 g/kg lactate (0.49 ± 0.07 vs 0.62 ± 0.04 s^-1^) groups. All other MTT comparisons were not statistically significant (all other p > 0.18).

TTP did not significantly change with lactate dose – with saline (5 ± 1 s), 0.5 g/kg lactate (7 ± 2 s), and 1 g/kg lactate (9 ± 4 s) (Group-wise p = 0.2) showing no statistically significant differences. All results shown in Figure 2.

**Figure 2.**
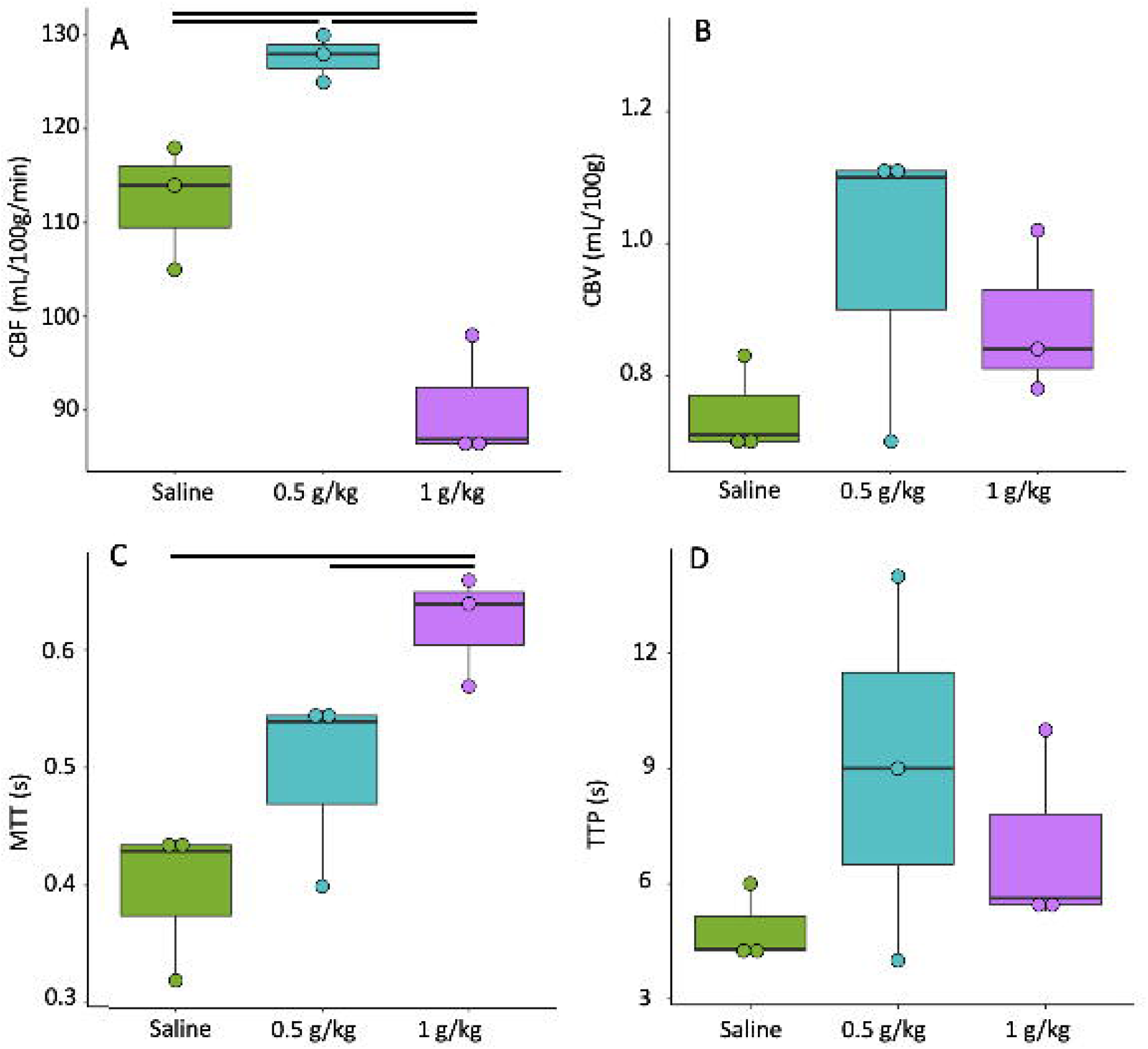
Results demonstrated significant differences between groups in Cerebral Blood Flow (A), and Mean Transit Time (B), but not Cerebral Blood Volume (C) or Time To Peak (D) between groups.

### Measurements of lactate exchange, but not PHD flux, vary with lactate dose

A significant difference in the [1-^13^C] lactate:[1-^13^C] pyruvate between the saline vs 0.5 g/kg lactate (0.22 ± 0.05 vs 0.59 ± 0.15) and the saline vs 1 g/kg lactate (0.22 ± 0.05 vs 0.62 ± 0.04) groups was observed, and this was accompanied by significant differences in the serum lactate at the end of scan between the saline and 0.5 g/kg lactate (1.3 ± 0.2 vs 2.6 ± 0.8 mmolL^-1^), the saline and 1 g/kg lactate (1.3 ± 0.2 vs 6 ± 2 mmolL^-1^) and the 0.5 g/kg and 1 g/kg lactate (2.6 ± 0.8 vs 6 ± 2 mmolL^-1^) groups.

There were no significant differences in ^13^C bicarbonate:[1-^13^C] pyruvate between any of the groups (Group-wise p = 0.1, 1 g/kg = 0.050 ± 0.007, 0.5 g/kg = 0.066 ± 0.014, Saline = 0.057±0.003), all results are shown in Figure 3.

**Figure 3.**
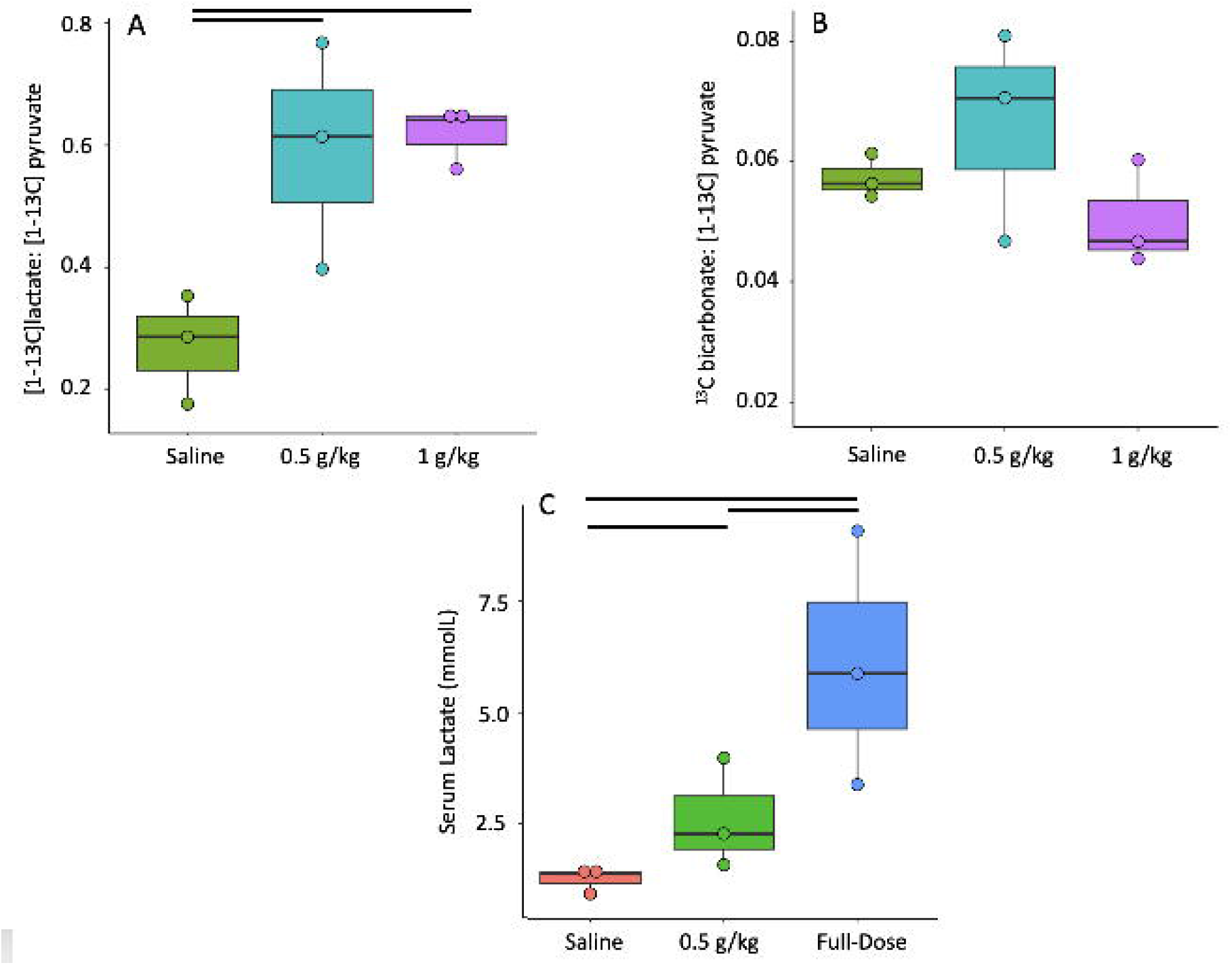
Results showed significant differences in [1-^13^C] lactate:[1-^13^C] pyruvate (A), serum lactate (B), but not measures ^13^C bicarbonate:[1-13C] pyruvate between groups.

### Sodium lactate administration leads to apparent uncoupling between cerebral perfusion and TCA metabolism

There were significant correlations between the [1-^13^C] lactate:[1-^13^C] pyruvate and serum lactate post scan (rho = 0.80) and between MTT and serum lactate post-scan (rho = 0.79). There was no significant correlation between ^13^C bicarbonate:[1-^13^C] pyruvate and CBF (p = 0.41). All statistically significant correlations are shown in figure 4.

**Figure 4.**
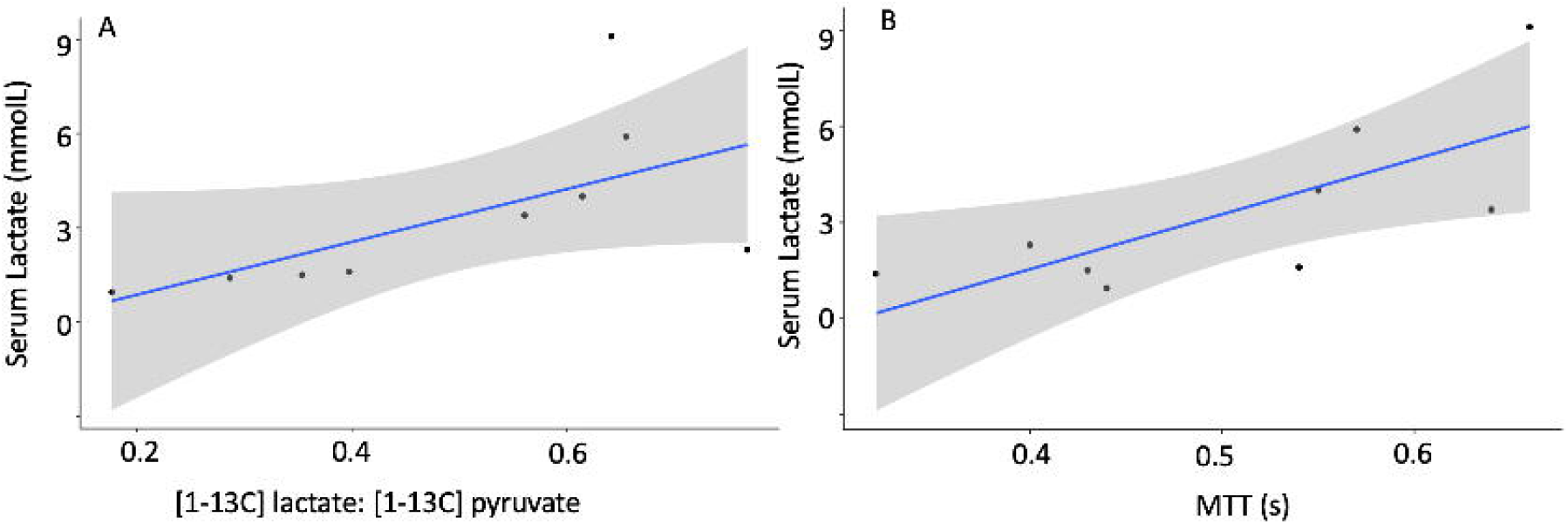
Significant correlations between [1-^13^C] lactate:[1-^13^C] pyruvate and serum lactate post scan (A, rho = 0.8) and MTT and serum lactate post scan (B, rho = 0.79)

## Discussion

This small study assessed the alterations in apparent metabolism and perfusion within the anesthetised rat brain due to administration of sodium lactate. Results indicated that measurements of cerebral perfusion were highly reproducible, providing increased confidence in the subsequent results assessing CBF post-lactate administration. Apparent measurements of lactate exchange and CBF in the anesthetised rodent brain appeared to be dependent on circulating serum lactate concentration. This modulation of apparent lactate exchange by adding non-labelled lactate is not a novel concept, having been demonstrated more than 20 years ago in cell-based experiments(18). However, the demonstration of this exchange phenomenon within the healthy rodent brain is novel and warrants further consideration for future pre-clinical and clinical studies. An indirect assessment of circulating lactate and apparent alterations in the [1-^13^C] lactate:[1-^13^C] pyruvate measured in the brain have been published, but not directly linked, in a previous study of a rodent model of cuprizone driven demyelination^23^. However, the authors of this study did also show that pyruvate dehydrogenase kinase activity was decreased in the rodent CNS, and so this may have also contributed to the change in [1-^13^C] lactate:[1-^13^C] pyruvate observed.

In this study it was noted that whilst [1-^13^C] lactate:[1-^13^C] pyruvate in the brain was susceptible to alterations in circulating lactate levels, measurements of ^13^C bicarbonate:[1-^13^C] pyruvate were unchanged, even with alterations in cerebral perfusion, suggesting that the [1-^13^C] lactate did not act as a major contributor to measurements of apparent PDH flux under these conditions. This may be because the [1-^13^C] pyruvate bolus in these experiments saturates PDH, and further dose dependency studies are required to answer this. These results could point toward an elevated oxygen extraction fraction within the brain compensating for reduced CBF, thus preserving apparent ^13^C bicarbonate production. This invariance of ^13^C bicarbonate with variations in CBF is encouraging, as it indicates that the study of cerebral PDH flux, and thus inferred TCA activity, and subsequent alterations due to pathologies such as stroke, multiple sclerosis, or traumatic brain injury, may be unaffected by variable tissue level serum lactate, thus providing an opportunity to understand injured mitochondria within the brain.

The results of this study suggest that an approach combining serum lactate, perfusion and ^13^C metabolic analysis with multi-variate regression may be needed to understand the [1-^13^C] lactate signal derived from studies of pathology, especially when looking at therapeutic response using hyperpolarized ^13^C MRI. A next step for research would be to use *in vivo* imaging to assess whether the change in [1-^13^C] lactate exchange is homogeneous across the whole brain. Finally, it is noted that the modulation of CBF by lactate is also not a new concept, with previous results showing an increase in CBF after low-dose administration, and this molecule shows promise for assisting in reperfusion post pre-clinical stroke^19,24,25^. The subsequent decrease in CBF due to high lactate has been previously observed in patients post stroke, and this may indicate a signalling pathway linking the two^26^.

### Limitations

The limitations of this study include the use of anesthetised animals, which do not fully reflect the human condition. Further studies to assess the validity of these results in a prospective human cohort with the acquisition of ^13^C imaging data would further clarify the extent of these results. Furthermore, the mechanism behind the control of CBF by lactate has not been elucidated in this study, with the potential for serum sodium, pH, blood pressure, or heart rate to be influencing CBF. Further work is needed to draw out the contribution of these individual components, and this study is by no means exhaustive – rather it highlights the challenges and potential pitfalls faced in the interpretation and acquisition of perfusion and metabolic imaging data. Indeed, one further contribution to the elevated [1-^13^C] lactate:[1-^13^C] pyruvate in the rodent brain may be due to [1-^13^C] lactate arriving from the blood in to the spectroscopy slice, not derived from cerebral tissue, however the spatial resolution of most imaging or spectroscopic methods, in the rodent brain, would be unable to separate out these two components due to a significant sequence point spread function, and diffusion weighted imaging may be hindered by low spatial resolution.

Finally, the results from this study are derived from a small number of animals. However, combining the reproducibility of CBF analysis, the large effect of lactate on CBF and metabolic measures, and the re-use of the same animals between each experiment to improves the statistical power of this study.

## Conclusion

This study has demonstrated the ability of sodium lactate to significantly alter CBF and the [1-^13^C]lactate signal, but to leave apparent PDH flux preserved in the anesthetised rodent brain. This has potential implications for the analysis and interpretation of future pre-clinical and clinical brain data using hyperpolarized ^13^C MRI.

## References

1. Venkat, P., Chopp, M. & Chen, J. New insights into coupling and uncoupling of cerebral blood flow and metabolism in the brain. Croatian Medical Journal 57, 223– 228 (2016).

2. Macvicar, B. A. & Newman, E. A. Astrocyte regulation of blood flow in the brain. Cold Spring Harbor Perspectives in Biology 7, 1–15 (2015).

3. Bélanger, M., Allaman, I. & Magistretti, P. J. Brain energy metabolism: Focus on Astrocyte-neuron metabolic cooperation. Cell Metabolism 14, 724–738 (2011).

4. Cheng, J. et al. Early glycolytic reprogramming controls microglial inflammatory activation. Journal of Neuroinflammation 18, 1–18 (2021).

5. Shivamurthy, V. K. N., Tahari, A. K., Marcus, C. & Subramaniam, R. M. Brain FDG PET and the diagnosis of dementia. AJR. American journal of roentgenology 204, W76– W85 (2015).

6. Brown, R. K. J., Bohnen, N. I., Wong, K. K., Minoshima, S. & Frey, K. A. Brain PET in Suspected Dementia: Patterns of Altered FDG Metabolism. RadioGraphics 34, 684– 701 (2014).

7. Sarikaya, I., Albatineh, A. N. & Sarikaya, A. Effect of various blood glucose levels on regional FDG uptake in the brain. Asia Oceania Journal of Nuclear Medicine and Biology 8, 46–53 (2020).

8. Bulte, D., Chiarelli, P., Wise, R. & Jezzard, P. Cerebral perfusion response to hyperoxia. Journal of cerebral blood flow and metabolismrz: official journal of the International Society of Cerebral Blood Flow and Metabolism 27, 69–75 (2007).

9. Wu, D. et al. The Spatiotemporal Evolution of MRI-Derived Oxygen Extraction Fraction and Perfusion in Ischemic Stroke. Frontiers in Neuroscience 15, 1–10 (2021).

10. Grist, J. T. et al. Hyperpolarized 13C MRI: A novel approach for probing cerebral metabolism in health and neurological disease. Journal of Cerebral Blood Flow and Metabolism (2020) xdoi:10.1177/0271678X20909045.

11. Le Page, L. M., Guglielmetti, C., Taglang, C. & Chaumeil, M. M. Imaging Brain Metabolism Using Hyperpolarized 13C Magnetic Resonance Spectroscopy. Trends in Neurosciences 43, 343–354 (2020).

12. Mammoli, D. et al. Kinetic Modeling of Hyperpolarized Carbon-13 Pyruvate Metabolism in the Human Brain. Transactions on Medical Imaging 0062, (2019).

13. Guglielmetti, C. et al. In vivo metabolic imaging of Traumatic Brain Injury. Scientific Reports 7, 17525 (2017).

14. Page, L. M. Le, Guglielmetti, C., Tiret, B. & Chaumeil, M. M. Hyperpolarized 13 C magnetic resonance spectroscopy detects toxin - induced neuroinflammation in mice. 1–11 (2019) doi:10.1002/nbm.4164.

15. Zaccagna, F. et al. Hyperpolarized carbon-13 magnetic resonance spectroscopic imaging: a clinical tool for studying tumour metabolism. The British Journal of Radiology 91, 20170688 (2018).

16. Bøgh, N. et al. Lactate saturation limits bicarbonate detection in hyperpolarized 13 C-pyruvate MRI of the brain. 1–10 (2022) doi:10.1002/mrm.29290.

17. Herzog, R. I. et al. Lactate preserves neuronal metabolism and function following antecedent recurrent hypoglycemia. Journal of Clinical Investigation 123, 1988–1998 (2013).

18. Berthet, C., Castillo, X., Magistretti, P. J. & Hirt, L. New evidence of neuroprotection by lactate after transient focal cerebral ischaemia: Extended benefit after intracerebroventricular injection and efficacy of intravenous administration. Cerebrovascular Diseases 34, 329–335 (2012).

19. Castillo, X. et al. A probable dual mode of action for both L-and D-lactate neuroprotection in cerebral ischemia. Journal of Cerebral Blood Flow and Metabolism 35, 1561–1569 (2015).

20. Bronikowski, T. A., Dawson, C. A. & Linehan, J. H. Model-free deconvolution techniques for estimating vascular transport functions. International Journal of Bio-Medical Computing 14, 411–429 (1983).

21. Koo, T. K. & Li, M. Y. A Guideline of Selecting and Reporting Intraclass Correlation Coefficients for Reliability Research. Journal of Chiropractic Medicine 15, 155–163 (2016).

22. Brindle, K. M., Campbell, I. D. & Simpson, R. J. A 1H n.m.r. study of the kinetic properties expressed by glyceraldehyde phosphate dehydrogenase in the intact human erythrocyte. Biochemical Journal 208, 583–592 (1982).

23. Guglielmetti, C. et al. Hyperpolarized 13C MR metabolic imaging can detect neuroinflammation in vivo in a multiple sclerosis murine model. Proceedings of the National Academy of Sciences of the United States of America 114, E6982–E6991 (2017).

24. Berthet, C., Castillo, X., Magistretti, P. J. & Hirt, L. New evidence of neuroprotection by lactate after transient focal cerebral ischaemia: Extended benefit after intracerebroventricular injection and efficacy of intravenous administration. Cerebrovascular Diseases 34, 329–335 (2012).

25. Buscemi, L. et al. Extended preclinical investigation of lactate for neuroprotection after ischemic stroke. Clinical and translational neuroscience 1, 1–9 (2020).

26. Rostami, E. et al. The correlation between cerebral blood flow measured by bedside xenon-CT and brain chemistry monitored by microdialysis in the acute phase following subarachnoid hemorrhage. Frontiers in Neurology 8, 1–7 (2017).

